# Piezo1 links cytoskeletal remodeling to differential YAP and β-catenin signaling in response to mechanical cues

**DOI:** 10.64898/2026.05.28.728382

**Authors:** Yuning Wu, Chang Ge, Zhi Hong Su, Lidan You, Fei Geng

## Abstract

Mechanical cues from the extracellular matrix regulate cancer cell behavior, but how these inputs are translated into distinct nuclear signaling responses remains incompletely understood. This study examined whether Piezo1, a mechanosensitive ion channel, contributes to cytoskeletal remodeling and differential regulation of YAP and β-catenin localization in breast cancer cells exposed to defined mechanical cues.

MDA-MB-231 breast cancer cells were cultured on substrates of defined stiffness and analyzed after Piezo1 knockdown or pharmacological modulation of Piezo1, Src signaling, myosin II activity, and actin polymerization. Nuclear localization of YAP and β-catenin was assessed by immunofluorescence imaging, cytoskeletal organization was evaluated using filamentous and globular actin staining, protein phosphorylation was analyzed by capillary electrophoresis-based immunoblotting, and cell migration was assessed using a wound-healing assay.

Piezo1 knockdown reduced YAP nuclear localization and increased β-catenin nuclear localization, while Piezo1 activation partially reversed these localization changes. Piezo1 knockdown also disrupted filamentous actin organization, and pharmacological disruption of actin polymerization produced similar effects on YAP and β-catenin localization. Piezo1 knockdown selectively reduced YAP tyrosine phosphorylation without altering canonical Hippo-associated YAP serine phosphorylation, and inhibition of Src signaling produced effects similar to Piezo1 knockdown. Functionally, Piezo1 knockdown impaired stiffness-dependent cell migration.

These findings support a role for Piezo1 in linking extracellular mechanical cues to cytoskeletal organization and differential regulation of YAP and β-catenin localization in breast cancer cells. This work provides a framework for understanding how mechanosensitive ion-channel signaling may contribute to context-dependent nuclear signaling responses during cancer cell mechanotransduction.

## Introduction

The tumor microenvironment (TME) is shaped by both biochemical and mechanical cues that directly regulate cancer cell behavior [1–3]. Among these, extracellular matrix (ECM) stiffness is a well-recognized regulator of tumor progression, impacting cell proliferation, invasion, and therapy resistance [4–6]. The ability of cells to sense and respond to these mechanical inputs, referred to as mechanotransduction, is therefore central to understanding tumor progression and metastatic potential [7].

Key downstream effectors of mechanotransduction include the transcriptional regulators Yes-associated protein (YAP) [8–10] and β-catenin [9,11]. YAP is highly sensitive to cytoskeletal tension and substrate rigidity, translocating to the nucleus in response to increased mechanical stress to drive proliferative and pro-survival gene expression [7]. In contrast, β-catenin, classically associated with Wnt signaling, has more recently been recognized as responsive to mechanical cues, though its behavior appears to differ from that of YAP [12,13]. However, it remains unclear whether a single upstream mechanosensitive regulator can coordinate cytoskeletal remodeling to differentially control multiple signaling pathways.

The actin cytoskeleton plays a central role in this process by integrating mechanical inputs and regulating nuclear access of transcription factors [14,15]. The balance between filamentous (F-actin) and globular (G-actin) states influences YAP localization, in part through interactions with scaffold proteins such as angiomotin (AMOT) [16], while also modulating β-catenin signaling through mechanisms that remain less well defined. Although cytoskeletal dynamics are known to govern mechanotransduction outcomes [17], the upstream sensors that coordinate these processes and direct pathway-specific responses are not fully understood.

Mechanosensitive ion channels have emerged as key candidates for translating physical forces into intracellular signals [18]. Among these, Piezo1 is a stretch-activated ion channel that responds to membrane tension by initiating downstream signaling events [19,20]. Piezo1 has been implicated in diverse cellular processes, including cytoskeletal remodeling [21], migration [22], and differentiation [23], and is increasingly recognized for its role in cancer progression [21]. However, whether Piezo1 functions as an upstream regulator that integrates mechanical inputs to differentially control downstream signaling pathways has not been established.

In our previous work, we demonstrated that ECM stiffness regulates a coordinated YAP–β-catenin signaling axis, revealing a compensatory interplay between these pathways in metastatic breast cancer cells [9,24,25]. However, the upstream mechanisms that govern this divergence and enable selective routing of mechanical signals remained undefined.

Here, we investigate whether Piezo1 acts as a mechanosensitive regulator that links extracellular stiffness to cytoskeletal remodeling and differential YAP and β-catenin localization. Our findings support a role for Piezo1 activation in enhancing F-actin organization, promoting YAP nuclear localization, and restricting β-catenin nuclear accumulation in response to mechanical cues. Conversely, Piezo1 knockdown disrupts cytoskeletal organization and induces opposing effects on these pathways. Together, our findings identify Piezo1 as a potential upstream regulator of mechanical signaling that differentially routes cytoskeleton-dependent inputs to nuclear signaling outputs, providing a mechanistic framework for how cells decode extracellular stiffness into divergent nuclear signaling programs.

## Materials and methods

### Cell culture and reagents

Human breast cancer MCF-7 and MDA-MB-231 cells were obtained from the American Type Culture Collection (ATCC, Manassas, VA, USA). Both MCF-7 and MDA-MB-231 cell lines were cultured in Dulbecco’s Modified Eagle Medium (DMEM; Thermo Fisher Scientific, NY, USA) supplemented with 10% fetal bovine serum (FBS; Thermo Fisher Scientific, NY, USA) at 37 °C in a humidified atmosphere with 5% CO₂. Cells were passaged at 70–80% confluence using 0.25% Trypsin-EDTA (Sigma-Aldrich, St. Louis, MO, USA).

Yoda1 was obtained from MedChemExpress (Monmouth Junction, NJ, USA). Dasatinib was purchased from Cell Signaling Technology (MA, USA). Blebbistatin was obtained from STEMCELL Technologies (Vancouver, Canada), and Latrunculin A (LatA) was purchased from Thermo Fisher (NY, USA). Yoda1 was used as a Piezo1 agonist, Dasatinib as a Src-family kinase inhibitor, Blebbistatin as a myosin II inhibitor, and Latrunculin A as an actin polymerization inhibitor.

### siRNA transfection

Prior to transfection, cells were cultured until they achieved 50% confluency. We utilized pre-designed siRNAs from the ON-TARGETplus SMARTpool collection targeting Piezo1, along with a non-targeting control siRNA (Dharmacon, Horizon Discovery, Lafayette, CO, USA). Transfection was carried out by introducing 25 nM of the respective siRNA along with DharmaFECT transfection reagent (Dharmacon, Horizon Discovery, Lafayette, CO, USA) to the cells, and this process was conducted following the guidelines provided by the manufacturer.

### RNA extraction and qRT-PCR

Total RNA was isolated from cells using the RNeasy Mini Kit (QIAGEN, Germantown, MD, USA), following the manufacturer’s instructions. RNA concentrations were measured using a multimode reader (Agilent BioTek, USA). For qRT-PCR, iTaq™ Universal SYBR Green One-Step Kit (Bio-Rad, Hercules, CA, USA) was used. The following specific human primers were employed:

Piezo1 (forward: 5′-CATCTTGGTGGTCTCCTCTGTCT-3′; reverse: 5′-CTGGCATCCACATCCCTCTCATC-3′)

GAPDH (forward: 5′-CTCCTGCACCACCAACTGCT-3′; reverse: 5′-GGGCCATCCACAGTCTCCTG-3′).

GAPDH served as an internal control for gene expression quantification. Gene expression levels were quantified using the 2^-ΔCT method. Technical duplicates were performed for each of three biological replicates.

### Immunofluorescence staining and image analysis

Cells were fixed with 3.7% formaldehyde for 15 minutes at room temperature, then permeabilized with 0.3% Triton X-100 in PBS for 25 minutes. To block non-specific binding sites, 3% BSA in PBS was used for incubation at 4℃ overnight.

The corresponding primary antibody(s) was applied next: mouse anti-human YAP antibody (SC-101199, Santa Cruz, USA) and rabbit anti-β-catenin antibody (Cell Signaling Technology, Danvers, MA, USA), or rabbit anti-Piezo1 antibody (Novus Biologicals, Centennial, CO, USA). Cells were incubated with the primary antibodies at 4°C overnight. Following this, cells were washed three times with 0.3% Triton X-100 in PBS and incubated with secondary antibodies: fluorescein isothiocyanate (FITC)-conjugated goat anti-mouse (Sigma-Aldrich, St. Louis, MO, USA) and/or Alexa Fluor™ 568-conjugated donkey anti-rabbit IgG (H+L) (Sigma-Aldrich, St. Louis, MO, USA), for 1 hour at room temperature in the dark. All primary and secondary antibodies were diluted in 3% BSA in PBS to achieve their working concentrations.

After the secondary antibody treatment, cells underwent three washes with PBST and were then incubated with DAPI (Sigma-Aldrich, St. Louis, MO, USA) for 10 minutes at room temperature, followed by three washes with PBS. The stained cells were imaged using a fluorescent microscope at a 20x magnification.

Fraction of cells exhibiting YAP/β-catenin nuclear localization was determined by dividing the number of cells showing nuclear localization of YAP (or β-catenin), based on signal intensity, by the total number of cells counted across all images from that given well.

### F-actin and G-actin staining

To evaluate the integrity of the actin cytoskeleton, cells underwent fixation with 3.7% formaldehyde for 15 minutes. Following fixation, they were washed twice with PBS and then permeabilized using 0.3% Triton X-100 for 25 minutes. Subsequent washes with PBS preceded a blocking step, where cells were incubated in 3% BSA dissolved in PBS at 4°C overnight. The staining procedure was carried out at room temperature in the following order, with PBS washes between each step. Cells were treated with deoxyribonuclease-I, conjugated with Alexa Fluor™ 594 (Thermo Fisher Scientific, Waltham, MA, USA), to label G-actin. Alexa Fluor™ 488 Phalloidin (Thermo Fisher Scientific, Waltham, MA, USA) was applied to stain F-actin. DAPI (Sigma-Aldrich, St. Louis, MO, USA) staining was performed for 10 minutes to visualize nuclei.

During staining, plates were covered with aluminum foil to protect from light. The quantification of F/G actin signal intensity ratio was determined by Image Prime software. Briefly, cells were masked according to the DAPI signal and the F-/G-actin signals. Nuclear and cytoplasmic regions (whole cell area subtracted by the nucleus area) were identified based on the masking. The mean integrated fluorescence intensity from the designated nuclear or cytoplasmic regions for each channel (GFP for F-actin, Cy5 for G-actin) was then computed to determine the F/G ratio from the objects. All cells captured from 3 wells (3 biological replicates per condition) were used in data quantification.

### Co-immunoprecipitation

Primary YAP antibody (Cell Signaling Technology, Danvers, MA, USA) was added to GammaG beads (Cytiva, Marlborough, MA, USA) and the mixture was placed on a rotator at 4°C for 24 hours to facilitate the conjugation of the antibodies to the beads. The following day, cell lysates containing 600 µg of protein were added to the antibody-bead complex and incubated with rotation for 2 hours at 4°C to allow the target protein (YAP) to bind to the bead-antibody complex. The YAP-antibody-bead complex was then collected by centrifugation. To dissociate the beads from the complex, a dye-free Laemmli buffer containing 4% SDS, 20% glycerol, 0.125M Tris-Cl, and 10% 2-mercaptoethanol (or DTT) was added. The supernatant, containing YAP and its associated proteins, was isolated and stored at -80°C for subsequent immunoblotting analysis.

### BCA assay and capillary-based electrophoresis (CE) immunoblotting

Harvested cell pellets were subjected to a wash with 1X PBS (Bio-Rad, Hercules, CA, USA). Subsequently, the cells were lysed in a buffer comprising 1% Triton X-100, 50 mM Tris-HCl (pH 8.0), 100 mM NaCl, and 1 mM EDTA, supplemented with 1X protease inhibitor (Roche, Basel, Switzerland). The protein concentration was determined using the Pierce™ BCA Protein Assay Kit (Thermo Fisher Scientific, Waltham, MA, USA) and a Tecan microplate reader (Infinite M200 Pro, Tecan, Männedorf, Switzerland).

To assess the expression levels of the target proteins in the cell lysates, a capillary electrophoresis-based immunoassay (ProteinSimple, San Jose, CA, USA) was conducted following the manufacturer’s protocol. Briefly, cell lysates were adjusted to a concentration of 0.4 μg/μL using lysis buffer and sample buffer, supplemented with 5X fluorescent master mix provided in the kit (ProteinSimple, San Jose, CA, USA). The cell lysate mixtures were subjected to heating at 95 °C for 5 minutes prior to loading.

Antibodies employed in this experiment included primary YAP antibody (Cell Signaling Technology, Danvers, MA, USA), primary PYAP antibody (Cell Signaling Technology, Danvers, MA, USA), primary β-catenin antibody (Cell Signaling Technology, Danvers, MA, USA), primary active-β-catenin antibody (Cell Signaling Technology, Danvers, MA, USA), and secondary anti-rabbit antibody (ProteinSimple, San Jose, CA, USA). Cell lysate mixtures, antibodies, and other necessary reagents were loaded into an assay plate (detection range: 12-230 kDa, provided in the kit) as per the manufacturer’s instructions. The CE immunoblot was run using default settings, and the results were provided by Compass for Simple Western software v6.1 (Biotechne, Minneapolis, MN, USA).

The peak area values were utilized to quantify the signal intensity of the proteins of interest. For protein intensity normalization, the values were normalized by the total detected protein in that capillary (ProteinSimple, San Jose, CA, USA).

### Wound healing assay

To assess the effect of Piezo1 on cell migration in response to substrate stiffness, a wound healing assay was performed using MDA-MB-231 cells cultured on collagen-coated PDMS substrates (0.2, 2, and 32 kPa). Cells were seeded into 6-well plates and grown to confluency to form a uniform monolayer. Linear wounds were introduced using a sterile 200-μL pipette tip to scratch across the center of each well.

Following wound generation, cells were immediately transfected with either control siRNA or Piezo1-targeting siRNA according to the manufacturer’s protocol. Phase-contrast images of the wounds were acquired at 0 hours and again at 48 hours post-transfection using an inverted microscope.

For quantification, the wound area at each time point was measured using ImageJ software. The wound healing rate was calculated as the percentage of wound area closure:

> % Closure = [(Area at 0 h – Area at 48 h) | Area at 0 h] × 100

At least three independent fields per well and three biological replicates per condition were analyzed. Statistical significance was determined using a two-tailed Student’s t-test or one-way ANOVA where applicable.

### Statistical analysis

Statistical analyses and graphs were performed and generated using GraphPad Prism. Data are presented as mean ± standard deviation (SD). The specific sample size (n), definition of biological replicates, and detailed quantitative methodologies for each experiment are provided in the respective figure legends. Differences between two groups were assessed using the unpaired Student’s t-test. For multiple group comparisons, one-way or two-way ANOVA was employed. In all tests, P < 0.05 was considered statistically significant. Unless otherwise noted, all experiments were performed at least in triplicate.

## Results

### Piezo1 coordinates stiffness-dependent nuclear localization of YAP and β-catenin through distinct cytoskeletal mechanisms

Substrate stiffness is a well-established mechanical cue that regulates mechanotransduction pathways in cancer cells. YAP is a key transcriptional regulator that responds to mechanical inputs by translocating into the nucleus [26], where it modulates cellular behavior related to cell proliferation, migration, and invasion. Given the established role of Piezo1 as a mechanosensitive ion channel and its potential to integrate physical signals from the microenvironment, we investigated how substrate stiffness and Piezo1 influence the nuclear localization of YAP and β-catenin in metastatic breast cancer cells.

To investigate the role of Piezo1 in mechanotransduction, we cultured MDA-MB-231 breast cancer cells on collagen-coated polydimethylsiloxane (PDMS) substrates of defined stiffness (0.2, 2, and 32 kPa) to model varying mechanical environments. Immunofluorescence analysis demonstrated that in control cells, YAP predominantly localized to the nucleus on stiff (32 kPa) substrates, consistent with mechanical activation, whereas on softer substrates (0.2 and 2 kPa), YAP displayed a mixed distribution, with both cytoplasmic and nuclear presence. Notably, knockdown of Piezo1 significantly diminished YAP nuclear localization on the 2 and 32 kPa substrates (Fig 1A, B).

**Fig 1.**
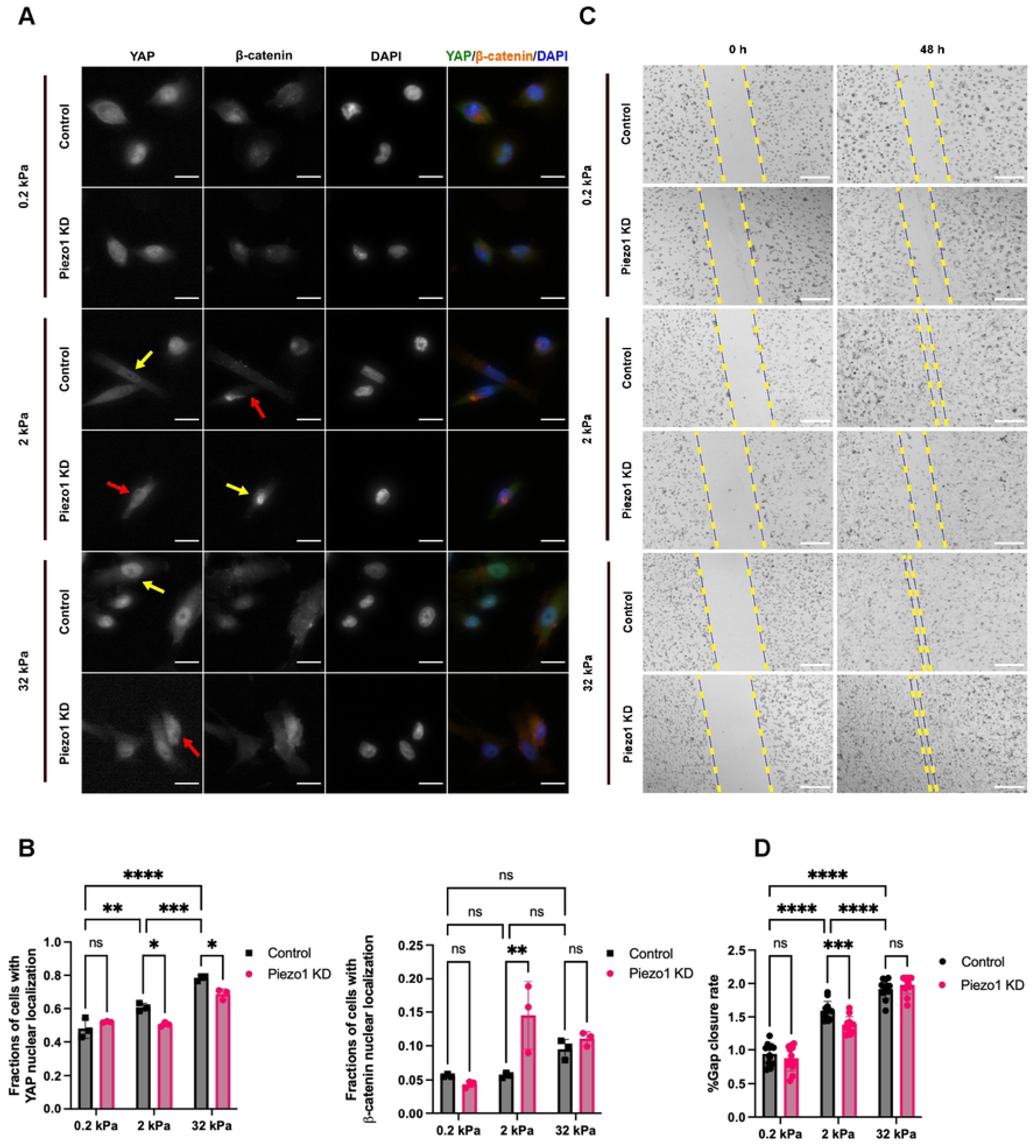
Piezo1 coordinates stiffness-dependent nuclear localization of YAP and β-catenin through distinct cytoskeletal mechanisms. (A) Representative IF staining of YAP and β-catenin. Cells were transfected with either non-target siRNA or Piezo1 siRNA for 48 hours. Scale bar = 20 µm. Yellow arrows indicate nuclear localization, whereas red arrows indicate cytoplasmic or non-nuclear localization. (B) Quantitative analysis of YAP (left) and β-catenin (right) localization results from the experiment in Fig 1A. For each biological replicate, multiple representative images were analyzed and the fraction of cells with nuclear localization was calculated. (C) Representative images from the wound healing assay over time for each treatment condition. The yellow dotted lines mark the wound boundary, and images are shown at 0 and 48 hours. Scale bars = 300 µm. (D) Quantification of wound closure percentage over time for each treatment condition. Data represent mean ± SD from n = 3 independent biological replicates. ns: not significant, *p < 0.05, **p < 0.01, ***p < 0.001, ****p < 0.0001.

In contrast, β-catenin nuclear localization was generally low and did not significantly vary across stiffness levels in control cells (Fig 1B, right). Notably, Piezo1 knockdown induced a slight but statistically significant increase in β-catenin nuclear localization at 2 kPa (p < 0.01), suggesting Piezo1 may modulate β-catenin localization under specific conditions.

To assess the role of substrate stiffness and Piezo1 activity in cell migration, we performed a wound healing assay on 0.2 kPa, 2 kPa and 32 kPa substrates and investigated the effect of Piezo1 knockdown on cell migratory potential (Fig 1C). Fig 1D shows that cells on the stiffer 32 kPa substrate close the wound more effectively than those on softer (2 and 0.2 kPa) substrates. In addition, quantitative analysis of wound closure percentage over time (Fig 1D) confirmed that control cells on the 2 kPa substrate exhibit significantly greater wound closure than the cells with Piezo1 knockdown, suggesting that Piezo1 supports efficient migration on 2 kPa surfaces.

These observations are consistent with our previous findings demonstrating that Piezo1 is required for stiffness-dependent cell migration and cellular behavior, supporting a functional role for Piezo1-mediated mechanotransduction in shaping cellular behavior.

### Mechanotransduction profiles differ between MDA-MB-231 and MCF-7 cells

We therefore compared Piezo1 expression, actin cytoskeletal organization, and mechanosensitive transcription factors between the highly metastatic MDA-MB-231 cells and the less metastatic MCF-7 cells.

The results highlight significant contrasts between these cell lines in terms of Piezo1 expression, actin filament organization, and levels of key mechanotransduction proteins, YAP and β-catenin. Immunofluorescence staining of Piezo1 and DAPI (Fig 2A, B) demonstrates a pronounced presence of Piezo1 in both MDA-MB-231 cells and MCF-7 cells.

**Fig 2.**
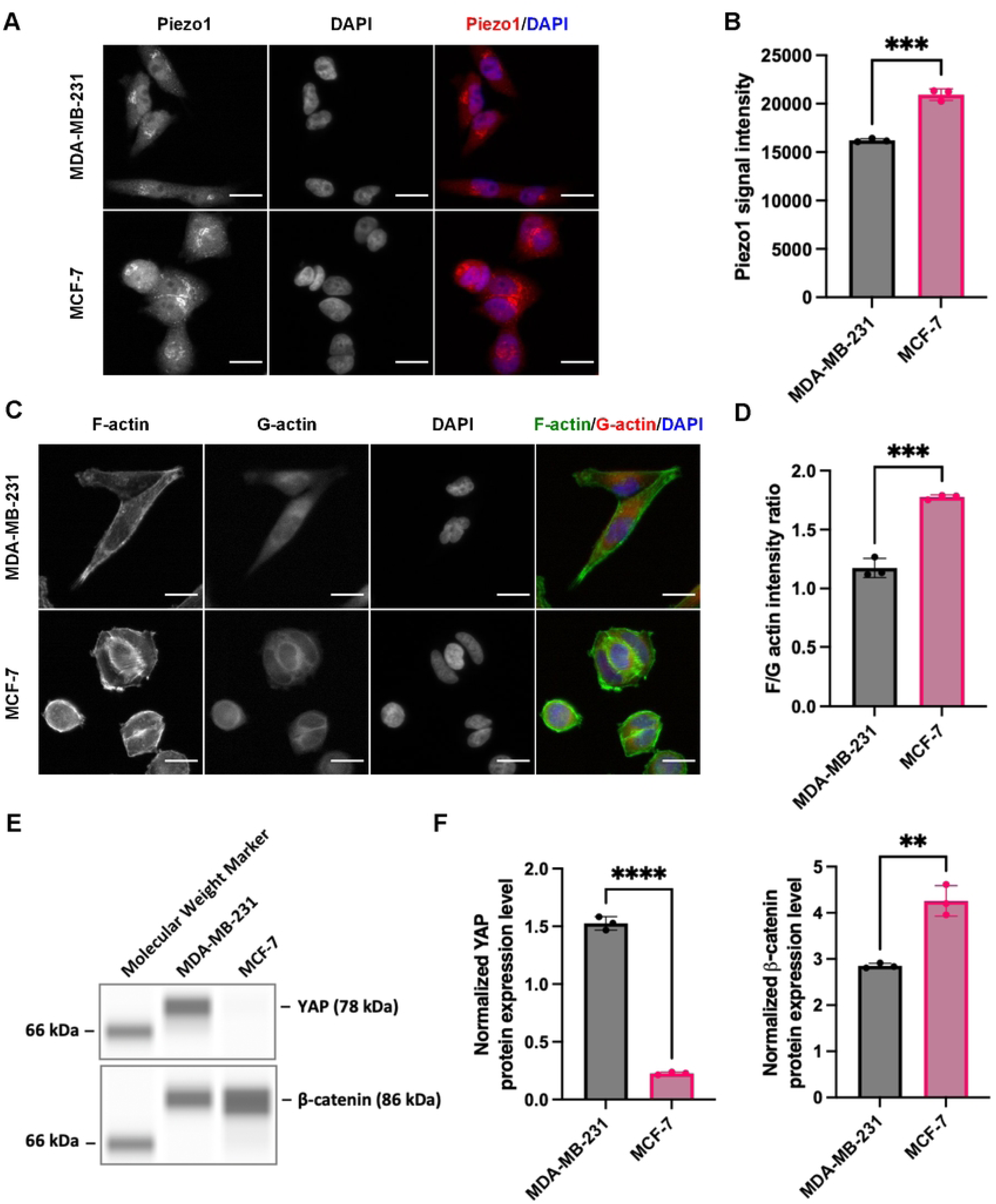
Mechanotransduction profiles differ between MDA-MB-231 and MCF-7 cells. (A) Immunofluorescence staining of Piezo1 (red) and nuclei (DAPI in blue) in MDA-MB-231 (top panel) and MCF-7 (bottom panel) cells. Scale bar = 20 µm. (B) Quantification of Piezo1 intensity in MDA-MB-231 and MCF-7 cells from the images in Fig 2A. (C) Immunofluorescence staining of F-actin (green), G-actin (red), and DAPI (blue) in MDA-MB-231 (top panel) and MCF-7 (bottom panel) cells. Scale bar = 20 µm. (D) Ratio of F-actin to G-actin in MCF-7 and MDA-MB-231 cells. (E) YAP and β-catenin protein expression in MCF-7 and MDA-MB-231 cells using quantitative capillary electrophoresis, highlighting differential expression levels of these proteins in the two cell lines. (F) Normalized YAP (left) and β-catenin (right) expression of Fig 2E in MDA-MB-231 and MCF-7 cells. Data represent mean ± SD from n = 3 independent biological replicates. **p < 0.01, ***p < 0.001, ****p < 0.0001.

The organization of the actin cytoskeleton, visualized through immunofluorescence staining of F-actin and G-actin (Fig 2C), shows that MCF-7 cells exhibit more structured F-actin filaments than MDA-MB-231 cells. This structural difference is quantified by the F-actin to G-actin ratio (Fig 2D), which is higher in MCF-7 cells. This higher F/G-actin ratio suggests enhanced cytoskeletal integrity and organization in MCF-7 [27]. Finally, quantitative capillary electrophoresis analysis of the mechanotransduction proteins YAP and β-catenin (Fig 2E) reveals that YAP protein levels are elevated in MDA-MB-231 cells compared to MCF-7 cells, while β-catenin shows higher expression in MCF-7 cells. This contrasting expression pattern suggests differential roles for YAP and β-catenin in mechanotransduction processes across the two cell lines. The expression values normalized to total protein (Fig 2F) further support that MDA-MB-231 cells exhibit a higher expression of YAP, which aligns with their enhanced mechanotransduction activity and metastatic potential.

### Piezo1 bidirectionally regulates YAP and β-catenin nuclear localization

To examine the specific role of Piezo1 in regulating YAP and β-catenin localization, we knocked down Piezo1 using siRNA in MDA-MB-231 cells and quantified nuclear versus cytoplasmic localization of YAP and β-catenin. qPCR confirmed efficient knockdown of Piezo1 mRNA levels compared to non-targeting controls (Fig 3A). Immunofluorescence staining revealed a marked shift in the intracellular distribution of YAP and β-catenin following Piezo1 knockdown: YAP was predominantly excluded from the nucleus, while β-catenin exhibited increased nuclear localization (Fig 3B). Quantitative analysis (Fig 3C) confirmed the observation, supporting a bidirectional regulatory effect.

**Fig 3.**
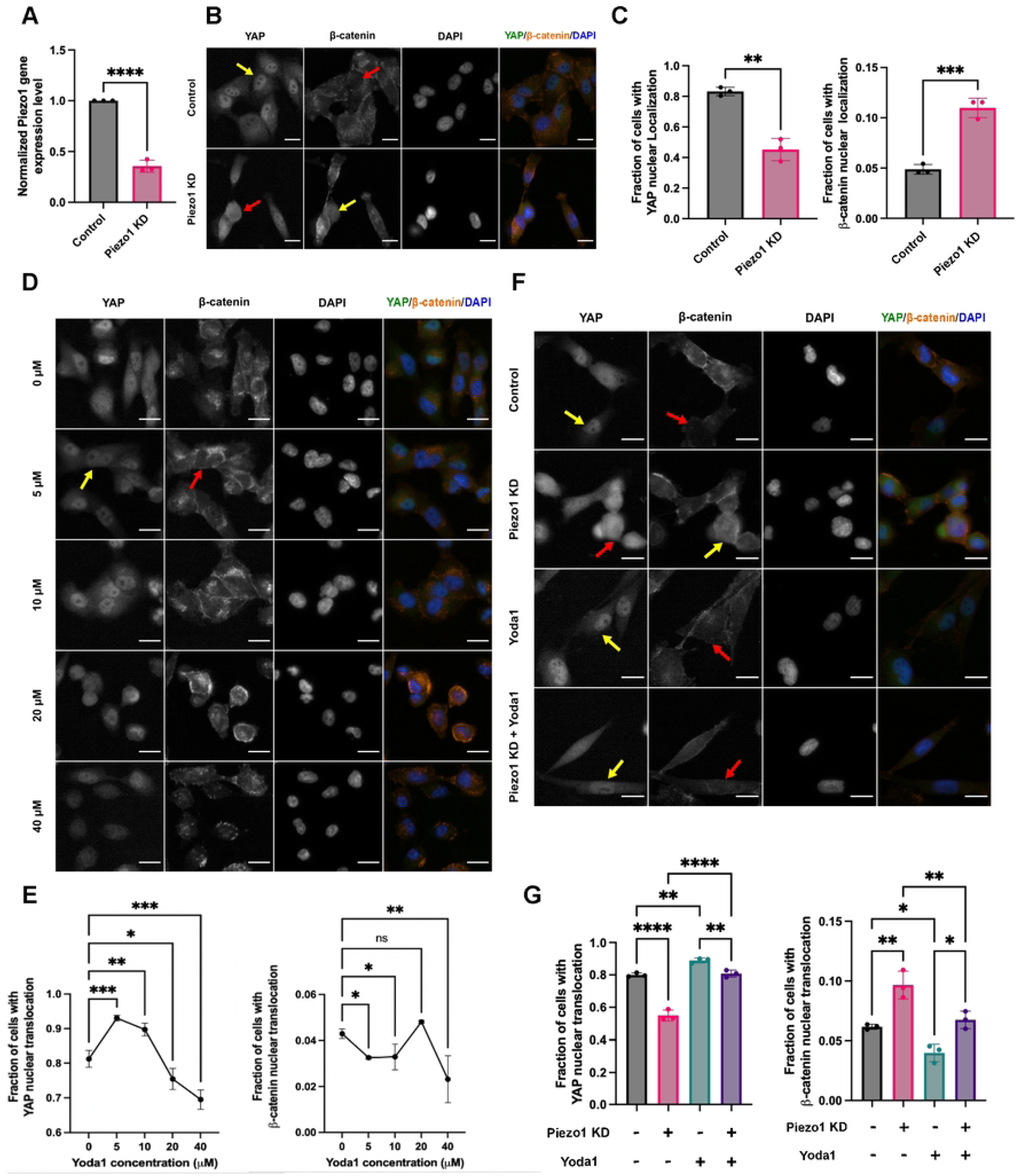
Piezo1 bidirectionally regulates YAP and β-catenin nuclear localization. (A) qRT-PCR showing Piezo1 knockdown efficiency 48 h post-transfection in MDA-MB-231 cells, normalized to GAPDH. (B) Representative immunofluorescence (IF) images of YAP and β-catenin localization after control or Piezo1 siRNA. Scale bar, 20 μm. Yellow arrows indicate nuclear localization, whereas red arrows indicate cytoplasmic or non-nuclear localization. (C) Quantification of nuclear YAP and β-catenin from (B). (D) IF images after 5 h treatment with increasing concentrations of Yoda1 or DMSO. (E) Quantification of nuclear YAP and â-catenin from (D). (F) IF images of cells treated with DMSO or 5 μM Yoda1 following Piezo1 knockdown. (G) Quantification of nuclear YAP and β-catenin from (F). For each biological replicate, multiple representative images were analyzed and the fraction of cells with nuclear localization was calculated. Data represent mean ± SD from n = 3 independent biological replicates. ns: not significant, *p < 0.05, **p < 0.01, ***p < 0.001, ****p < 0.0001.

Next, we treated cells with increasing concentrations of Yoda1 for 5 hours and observed a concentration-dependent restoration of YAP nuclear localization and suppression of β-catenin nuclear localization (Fig 3D, E). This effect was further validated in cells pre-treated with Piezo1 siRNA: Yoda1 treatment partially reversed the knockdown-induced mislocalization of both YAP and β-catenin (Fig 3F, G).

### Piezo1 regulates YAP tyrosine phosphorylation

Given our findings that Piezo1 knockdown disrupts YAP nuclear localization (Fig 3), we next examined whether Piezo1 influences YAP post-translational modifications, specifically phosphorylation at tyrosine 357 (Y357) and serine 127 (S127)—two sites known to regulate YAP localization and transcriptional activity. Capillary electrophoresis analysis revealed that total YAP protein levels remained unchanged following Piezo1 knockdown (Fig 4B). However, phosphorylation at Y357 was significantly reduced in Piezo1-depleted cells compared to controls (Fig 4D), while phosphorylation at S127 was unaffected (Fig 4C). These results suggest that Piezo1 selectively modulates YAP activity by regulating its tyrosine, but not serine, phosphorylation.

**Fig 4.**
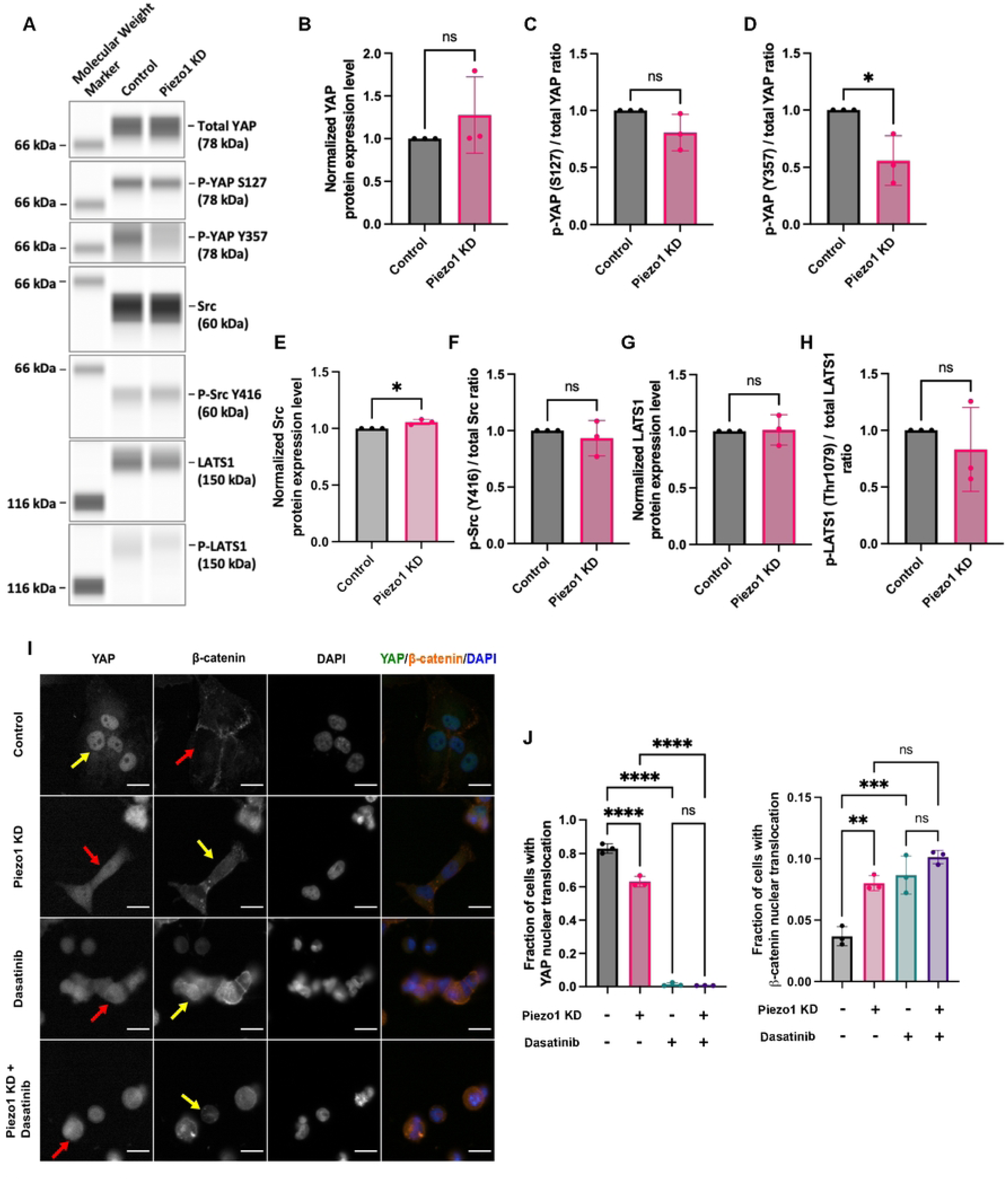
Src inhibition mimics Piezo1 knockdown effects on YAP and β-catenin. (A) Capillary electrophoresis-based immunoblot analysis of total and phosphorylated YAP, Src, and LATS1 at 48 hours following siRNA transfection. Groups: Control (non-target siRNA) and Piezo1 KD (Piezo1 siRNA). (B-H) Quantification from Fig 4A, showing fold changes in signal intensity: total YAP (B), phosphorylated S127 YAP (C), phosphorylated Y357 YAP (D), total Src (E), phosphorylated Src at Y416 as a percentage of total Src (F), total LATS1 (G), and p-LATS1 at Thr1079 as a percentage of total LATS1 (H). Capillary Western blot analysis was performed, and protein band intensities were quantified using Compass for SW software. All data were normalized to total protein levels and shown as fold changes relative to control. Phosphorylated protein levels are shown as fold changes relative to their non-phosphorylated forms. Statistical significance was determined by an unpaired, two-tailed Student’s t-test. p < 0.05; ns, not significant. (I) Representative IF staining of YAP and β-catenin. Cells were transfected with either non-target siRNA or Piezo1 siRNA for 48 hours. At 43 hours (5 hours before fixation), cells were either treated with 0.1 µM Dasatinib or an equivalent volume of DMSO. Scale bar = 20 µm. Red arrows: nuclear exclusion; yellow arrows: nuclear localization. (J) Quantitative analysis of the cell fraction with the nuclear localization of YAP (left) and β-catenin (right) localization results from experiment (I). Data represent mean ± SD. n = 3 independent biological replicates (wells). For each well, the fraction of cells with nuclear localization was quantified from multiple representative images by dividing the number of cells with YAP/β-catenin nuclear localization by the total number of cells. Statistical significance was assessed using one-way ANOVA followed by Tukey’s post hoc test. ns: not significant, **p < 0.01, ***p < 0.001, ****p < 0.0001.

### Src inhibition mimics Piezo1 knockdown effects on YAP and β-catenin

Since YAP phosphorylation at Y357 is mediated by Src [28,29], we investigated the effects of Src inhibition on the localization of YAP and β-catenin, as well as whether Src activity was influenced by Piezo1 knockdown. Firstly, we analyzed the YAP/β-catenin nuclear localization in cells transfected with control (non-targeting siRNA) or Piezo1 siRNAs, and/or treated with Dasatinib (a Src inhibitor). In control cells, YAP showed nuclear localization, as indicated by yellow arrows, while Piezo1 knockdown significantly reduced YAP nuclear localization, with more prominent cytoplasmic localization (red arrows) (Fig 4I).

Quantitative analysis confirmed a significant decrease in the proportion of cells with nuclear YAP in both the Piezo1 knockdown and Dasatinib-treated groups (Fig 4J, left). Interestingly, β-catenin nuclear localization was enhanced under both conditions (Fig 4J, right), consistent with previous findings (Fig 3).

Since Src inhibition significantly reduced YAP nuclear localization, we next examined whether Piezo1 knockdown alters Src and LATS1 activity—two well-established upstream regulators of YAP that act via distinct phosphorylation mechanisms: Src phosphorylates YAP at Y357 to promote its nuclear localization, while LATS1 phosphorylates YAP at S127 to retain it in the cytoplasm [30,31]. Capillary electrophoresis immunoblotting showed a significant increase in total Src protein levels following Piezo1 knockdown (Fig 4E). However, phosphorylation at Src tyrosine 416 (pSrc-Y416), a marker of Src activation [32], remained unchanged (Fig 4F) and may reflect altered spatial or cytoskeleton-associated Src signaling. These findings suggest that Piezo1-dependent cytoskeletal remodeling may influence YAP localization through Src-associated signaling [33].

Similarly, total LATS1 levels and its phosphorylation at threonine 1079 (p-LATS1), a marker of its activation [34], remained unchanged (Fig 4G, H). These findings suggest that the reduction in YAP nuclear localization following Piezo1 knockdown could be regulated through Src-dependent signaling rather than changes in Src activation levels.

### Myosin II and actin dynamics differentially influence YAP and β-catenin localization

The cytoskeleton has been shown to play a pivotal role in Piezo1 and YAP signaling in cancer [35,36]. To further investigate Piezo1 regulation on YAP and β-catenin nuclear localization, we examined how myosin and actin filaments impact Piezo1 activity. Specifically, we explored the effects of myosin inhibition on YAP and β-catenin localization by treating cells with blebbistatin, a myosin inhibitor [37]. Interestingly, blebbistatin treatment alone did not significantly impact YAP nuclear localization compared to the control, suggesting that myosin activity is not directly required for YAP’s nuclear localization. However, when blebbistatin was combined with Piezo1 knockdown, a notable increase in YAP nuclear localization was observed (Fig 5A, B), highlighting a potential compensatory mechanism where alternative pathways may promote YAP nuclear entry in the absence of Piezo1. For β-catenin, blebbistatin treatment alone led to an increase in nuclear localization compared to the control, suggesting that myosin inhibition enhances β-catenin’s nuclear localization.

**Fig 5.**
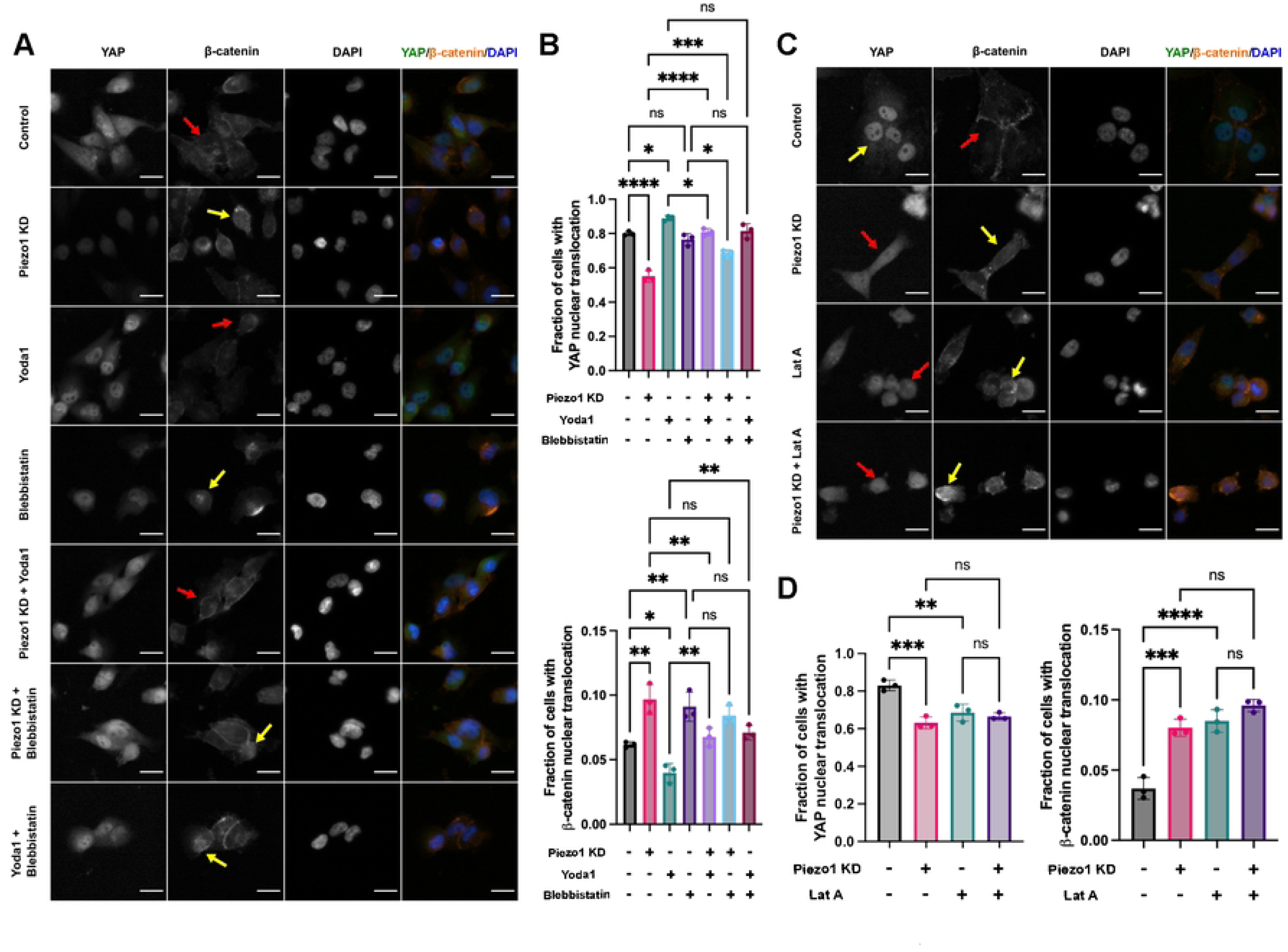
Myosin II and actin dynamics differentially influence YAP and β-catenin localization. (A) The effect of Blebbistatin treatment on Piezo1-mediated YAP and β-catenin nuclear localization. Cells were transfected with either non-target siRNA or Piezo1 siRNA for 48 hours before fixation, staining, and imaging. At 40 hours, cells were either treated with drugs (5 μM Yoda1, 5 μM Blebbistatin, or combination of 5 μM Yoda1 and 5 μM Blebbistatin) or an equivalent volume of DMSO. Scale bar = 20 µm. Yellow arrows indicate nuclear localization, whereas red arrows indicate cytoplasmic or non-nuclear localization. (B) Quantitative analysis of protein localization results from experiment in Fig 5A. (C) The effect of LatA treatment on Piezo1-mediated YAP and β-catenin nuclear localization. Cells were transfected with either non-target siRNA or Piezo1 siRNA for 48 hours before fixation, staining, and imaging. At 40 hours (5 hours before fixation), cells were either treated with drugs (5 μM Yoda1, 0.1 μM LatA, or combination of 5 μM Yoda1 and 0.1 μM LatA) or an equivalent volume of DMSO. Scale bar = 20 µm. (D) Quantitative analysis of protein localization results from experiment in Fig 5C. For each biological replicate, multiple representative images were analyzed and the fraction of cells with nuclear localization was calculated. Data represent mean ± SD from n = 3 independent biological replicates. ns: not significant, *p < 0.05, **p < 0.01, ***p < 0.001, ****p < 0.0001.

We further investigated the role of the actin cytoskeleton in YAP and β-catenin localization by treating cells with LatA, which disrupts actin polymerization. LatA treatment alone resulted in reduced YAP nuclear localization, indicating that an intact actin cytoskeleton supports YAP’s nuclear entry. When LatA was combined with Piezo1 knockdown, no additional reduction in YAP nuclear localization was observed compared to Piezo1 knockdown alone (Fig 5C, D), suggesting that Piezo1 may play a more dominant role or that its loss saturates the effects of actin disruption on YAP translocation. For β-catenin, LatA treatment alone led to an increase in nuclear localization (Fig 5C, D), implying that actin depolymerization enhances β-catenin nuclear entry.

Importantly, treatment with Yoda1, a Piezo1-specific agonist, partially reversed the effects of blebbistatin. Yoda1 restored YAP nuclear localization in conditions where it was suppressed by myosin inhibition (Fig 5A, B), supporting the role of Piezo1 activation in promoting cytoskeletal integrity. For β-catenin, Yoda1 treatment reduced nuclear localization induced by blebbistatin, demonstrating that Piezo1 activation can antagonize the β-catenin nuclear accumulation triggered by cytoskeletal disruption.

### Piezo1 promotes F-actin assembly to support YAP and β-catenin nuclear localization

To dissect the role of Piezo1 in cytoskeletal remodeling and its impact on YAP and β-catenin nuclear localization, we assessed F-actin and G-actin organization in MDA-MB-231 cells following Piezo1 knockdown or treatment with various cytoskeletal modulators. Cells were exposed to Yoda1 (Piezo1 agonist), LatA (an actin polymerization inhibitor), Blebbistatin (a myosin II inhibitor), and Dasatinib (a Src inhibitor).

Immunofluorescence analysis revealed that Yoda1 treatment enhanced F-actin intensity in control cells (yellow arrows, Fig 6A), indicating that Piezo1 activation promotes actin polymerization. In contrast, Piezo1 knockdown led to a marked reduction in F-actin signal (red arrows, Fig 6A), implicating Piezo1 as a key regulator of actin cytoskeletal integrity. As expected, LatA substantially disrupted F-actin architecture. Blebbistatin induced less pronounced changes in F-actin structure, suggesting that Piezo1 may act in coordination with myosin II and Src to maintain cytoskeletal homeostasis. Quantitative analysis of F-actin and G-actin staining intensities (Fig 6B) confirmed that both Piezo1 knockdown and LatA treatment significantly diminished F-actin levels, as determined by one-way ANOVA.

**Fig 6.**
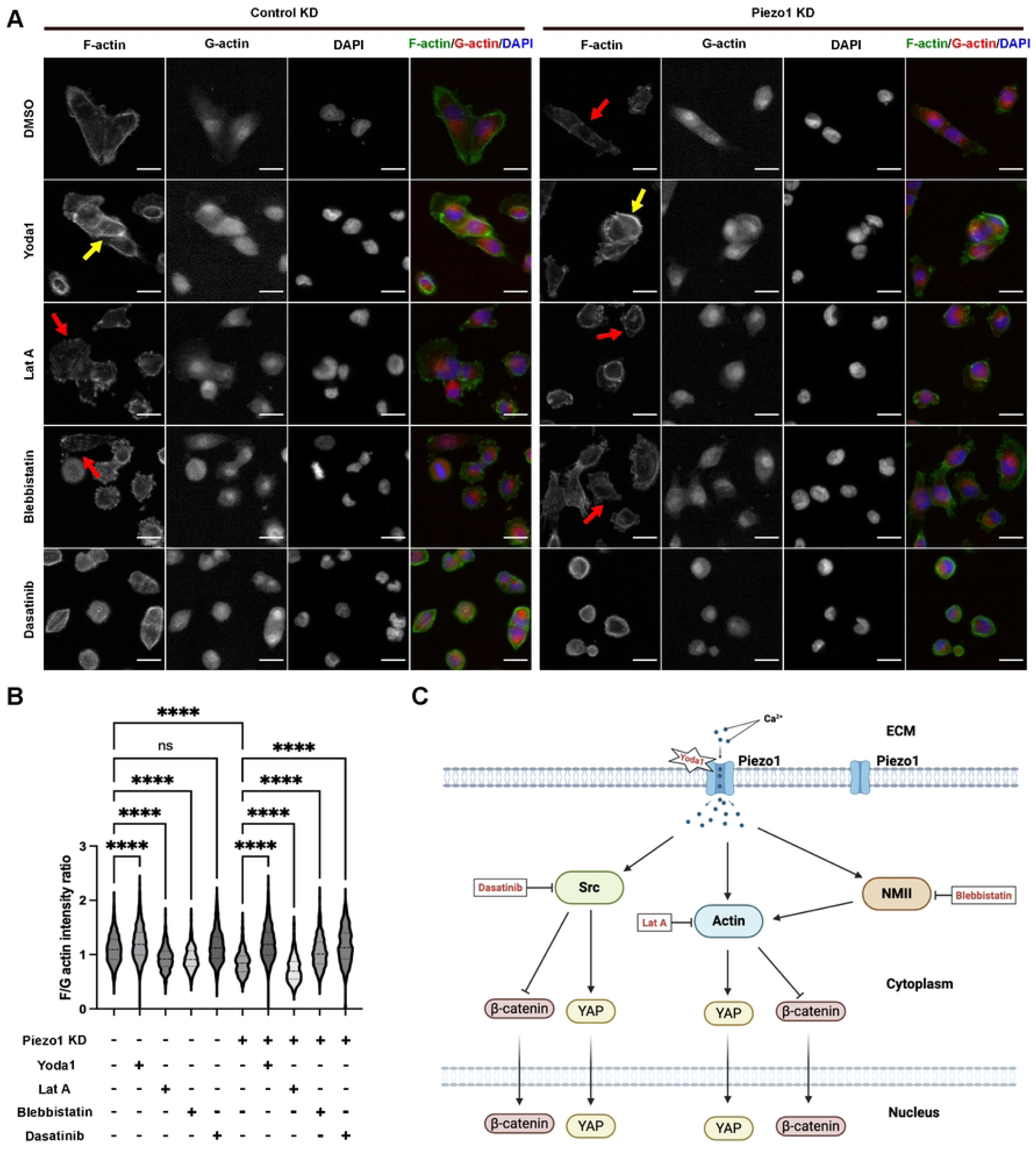
Piezo1 promotes F-actin assembly to support YAP and β-catenin nuclear localization. (A) Representative F- and G-actin staining in MDA-MB-231 cells transfected with control or Piezo1 siRNA for 48 h, followed by treatment with DMSO, Yoda1 (5 μM), LatA (0.1 μM), Blebbistatin (5 μM), Dasatinib (0.1 μM), or Yoda1 + Blebbistatin. Scale bar, 20 μm. Arrows indicate representative regions of altered F-actin organization. (B) Quantification of F- and G-actin intensity across conditions, shown as violin plots. Data represent mean ± SD from n = 3 independent biological replicates. (C) Working model: Piezo1 activation is proposed to engage NMII to promote F-actin assembly, thereby potentially differentially regulating YAP and β-catenin nuclear localization. Inhibitors disrupt this pathway by altering cytoskeletal dynamics. ns: not significant, ****p < 0.0001.

Based on these results, we propose a working model (Fig 6C) in which mechanical activation of Piezo1 engages downstream effectors such as NMII to promote F-actin polymerization. The assembled F-actin cytoskeleton may then serve as a scaffold that differentially regulates the nuclear localization of YAP and β-catenin.

### Piezo1-dependent cytoskeletal remodeling regulates YAP localization

Given the established role of AMOT in connecting the cytoskeleton to YAP regulation [38,39] and that F-actin and YAP compete for binding to AMOT [40], we next sought to characterize YAP-AMOT interaction and F-actin dynamics in the cells with Piezo1 knockdown. Fig 7A demonstrates that YAP was immunoprecipitated using an anti-YAP antibody, and AMOT was detected in the same complex, confirming the interaction between YAP and AMOT in MDA-MB-231 cells.

**Fig 7.**
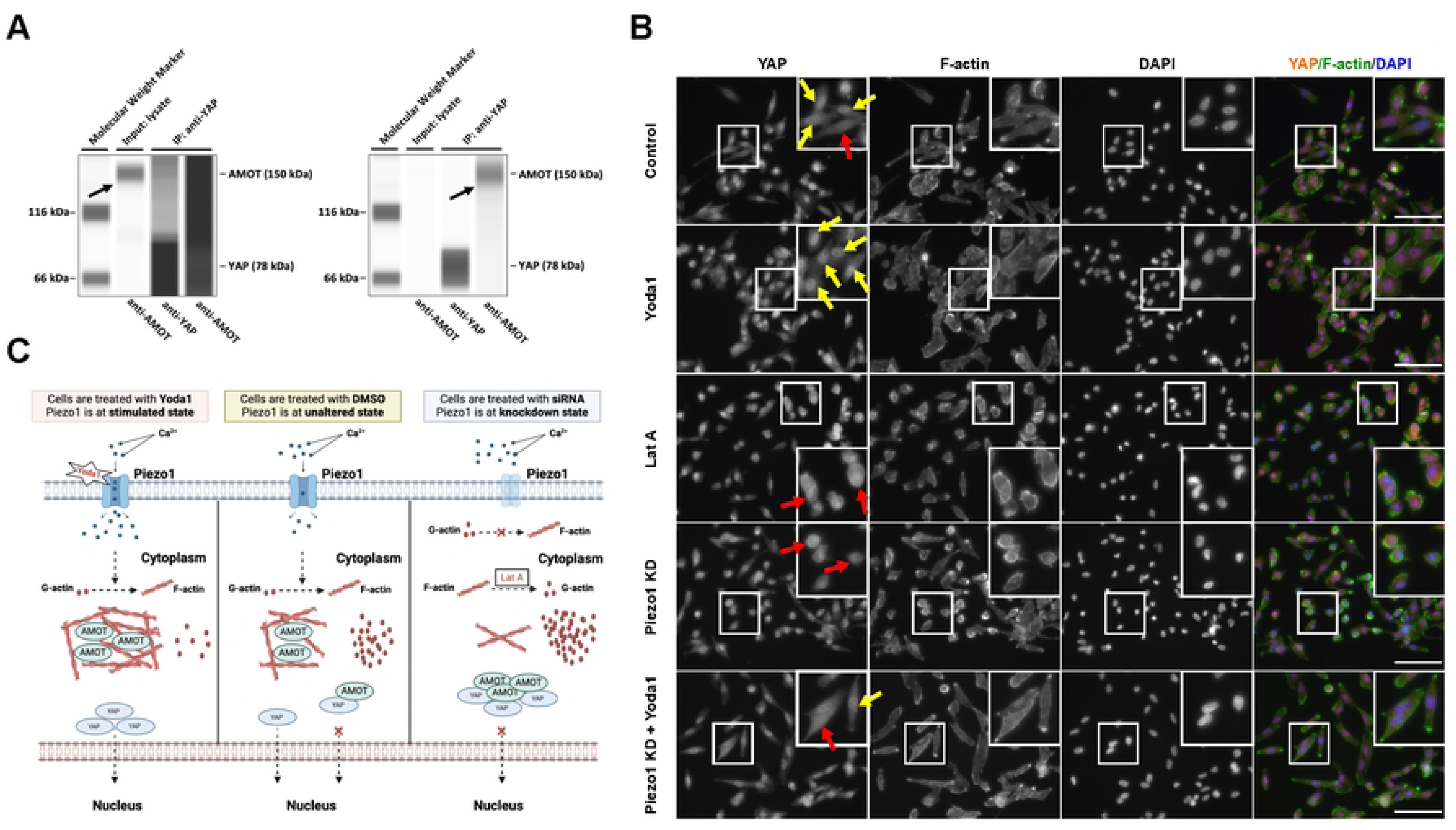
Piezo1-dependent cytoskeletal remodeling regulates YAP localization. (A) Capillary electrophoresis immunoblots showing YAP–AMOT interaction. The left panel shows a longer exposure blot, allowing better visualization of AMOT expression in the input lysate. The right panel shows a regular exposure blot of the immunoprecipitation experiment, where YAP was pulled down in the IP sample using an anti-YAP antibody, and co-immunoprecipitated AMOT was detected using an anti-AMOT antibody. (B) Representative immunofluorescence images of YAP (red), F-actin (green), and nuclei (DAPI, blue) in control, Yoda1-treated, LatA-treated, Piezo1 KD, and Piezo1 KD + Yoda1 groups. Yoda1 enhances YAP nuclear localization and F-actin organization, while Piezo1 KD and LatA disrupt both. Magnified views show YAP–F-actin colocalization. Scale bar, 10 μm. (C) Proposed model: Piezo1 loss or actin disruption leads to increased YAP cytoplasmic retention.

Given that YAP localization is influenced by both actin cytoskeletal organization and AMOT-associated regulation [40], we further examined YAP and F-actin dynamics in the cells with Piezo1 knockdown or activation (Yoda1 treatment). Fig 7B shows that in control cells, a majority of cells exhibit YAP localization in the nucleus, with intact F-actin structures supporting its nuclear localization. Yoda1 treatment enhances YAP nuclear localization and promotes colocalization with F-actin, indicating that Piezo1 activation facilitates YAP nuclear entry by stabilizing F-actin structures. In contrast, Piezo1 knockdown disrupts F-actin organization, reducing YAP nuclear localization and emphasizing the critical role of Piezo1 in maintaining the balance between F-actin polymerization and YAP nuclear localization. Similarly, LatA treatment reduces YAP nuclear localization by disrupting actin polymerization, further underscoring the dependence of YAP nuclear entry on intact F-actin.

We propose a model in which Piezo1-dependent F-actin organization regulates YAP localization through cytoskeleton-dependent mechanisms (Fig 7C). Under basal conditions, Piezo1 activity supports F-actin polymerization and is correlated with efficient YAP nuclear localization. Piezo1 activation (Yoda1 treatment) enhances F-actin assembly, which correlates with reduced YAP cytoplasmic retention and increased nuclear localization (Fig 7C). In contrast, Piezo1 downregulation disrupts F-actin organization and reduces YAP nuclear localization, leading to increased cytoplasmic retention. Similarly, actin depolymerization by LatA impairs F-actin integrity and reduces YAP nuclear localization (Fig 7C).

## Discussion

Mechanotransduction enables cells to convert mechanical cues from the extracellular environment into intracellular signaling responses that regulate cellular behavior [41,42]. In cancer, increased ECM stiffness is coupled with enhanced proliferation, migration, and metastatic potential [43]. While transcriptional regulators such as YAP and β-catenin have been widely implicated as downstream effectors of mechanical signaling, how cells differentially regulate these pathways in response to mechanical inputs remains incompletely understood [44]. In this study, we identify Piezo1 as a mechanosensitive regulator that drives cytoskeletal remodeling and differentially influences the nuclear localization of YAP and β-catenin (Figs 1 and 6).

We first show that Piezo1 regulates stiffness-dependent YAP nuclear localization while exerting an opposing effect on β-catenin (Fig 1). Under increasing substrate stiffness, YAP nuclear localization is enhanced, whereas Piezo1 knockdown disrupts this response (Fig 1A, B). In contrast, β-catenin nuclear localization is modest under basal conditions but shows a slight increase upon Piezo1 knockdown at intermediate stiffness (Fig 1B). These observations suggest that Piezo1 differentially regulates YAP and β-catenin localization under specific mechanical conditions. Functionally, Piezo1 knockdown impairs stiffness-dependent cell migration (Fig 1C, D), supporting a role for Piezo1-mediated mechanotransduction in regulating cellular behavior.

Our data further support a link between Piezo1 activity and cytoskeletal organization. Comparative analysis between MDA-MB-231 and MCF-7 cells reveals differences in Piezo1 expression, actin architecture, and YAP/β-catenin expression profiles (Fig 2). Notably, variations in F-actin organization correlate with differences in mechanosensitive signaling proteins (Fig 2C, D), suggesting that cytoskeletal state reflects distinct signaling contexts. Consistent with this, direct manipulation of Piezo1 demonstrates that Piezo1 knockdown reduces YAP nuclear localization and increases β-catenin nuclear accumulation, while pharmacological activation using Yoda1 partially reverses these effects (Fig 3G). Together, these findings indicate that Piezo1 activity bidirectionally regulates these pathways.

Although mechanistic experiments were primarily conducted in MDA-MB-231 cells, comparative analyses with MCF-7 cells support the broader relevance of Piezo1-associated mechanotransduction in breast cancer cell behavior.

Mechanistically, our results suggest that Piezo1 regulates YAP localization through a cytoskeleton-associated, Hippo-independent pathway. Piezo1 knockdown selectively reduces phosphorylation of YAP at Y357 without affecting S127 phosphorylation (Fig 4), indicating that canonical Hippo signaling is not the primary regulator in this context. Inhibition of Src signaling phenocopies the effects of Piezo1 knockdown, reducing YAP nuclear localization and increasing β-catenin nuclear accumulation (Fig 4I, J). However, while total Src levels increase following Piezo1 knockdown, Src activation remains unchanged (Fig 4E, F), suggesting involvement of Src-associated signaling rather than global changes in Src activity.

The actin cytoskeleton emerges as an important mediator of these effects. Disruption of actin polymerization using Latrunculin A reduces YAP nuclear localization and increases β-catenin nuclear accumulation, consistent with the effects observed following Piezo1 knockdown (Fig 5C, D). In contrast, myosin inhibition produces more variable effects (Fig 5A, B), suggesting that actin polymerization may play a more prominent role than contractility in regulating nuclear localization in this system. Importantly, Piezo1 activation partially rescues these perturbations, supporting a role for Piezo1 in maintaining cytoskeletal organization.

Direct visualization of cytoskeletal architecture confirms that Piezo1 activity drives cytoskeletal remodeling. Piezo1 activation enhances F-actin assembly, whereas Piezo1 knockdown or pharmacological disruption reduces F-actin levels (Fig 6A, B). These findings support a model in which Piezo1 activity orchestrates cytoskeletal remodeling, with F-actin acting as an intermediary influencing YAP and β-catenin localization (Fig 6C).

At the molecular level, our findings suggest that Piezo1-dependent cytoskeletal organization may influence YAP localization through AMOT-associated mechanisms. Co-immunoprecipitation confirms interaction between YAP and AMOT (Fig 7A), and imaging analysis shows that Piezo1 activation enhances F-actin organization and YAP nuclear localization, whereas Piezo1 knockdown or actin disruption leads to cytoplasmic retention of YAP (Fig 7B). These observations are consistent with a model in which cytoskeletal organization influences YAP localization through competition with AMOT-mediated sequestration (Fig 7C).

In contrast to YAP, β-catenin exhibits a distinct response to mechanical and cytoskeletal perturbations [45,46]. Increased β-catenin nuclear localization following Piezo1 knockdown or actin disruption (Figs 1, 3, and 5) is consistent with cytoskeleton-sensitive regulation of β-catenin localization. Notably, the magnitude of this effect is modest and appears dependent on specific mechanical conditions (Fig 1B), suggesting that β-catenin may respond to cytoskeletal changes differently than YAP. Although YAP and β-catenin exhibit opposing localization patterns in this study, further work will be required to determine whether these responses are mechanistically coupled or independently regulated downstream of Piezo1.

From a broader perspective, these findings support a model in which Piezo1 functions as a mechanosensor and a signaling node that selectively routes cytoskeleton-dependent mechanical inputs toward distinct nuclear signaling outcomes. Consistent with its effects on cytoskeletal organization, Piezo1 promotes YAP nuclear localization and restricts β-catenin nuclear accumulation (Figs 3 and 6). This differential regulation may provide a mechanism by which cells respond to mechanical cues in a context-dependent manner.

While our findings support a role for Piezo1 in modulating YAP tyrosine phosphorylation through Src-associated signaling, the precise molecular intermediates linking Piezo1 activation to cytoskeleton-associated Src signaling remain to be fully defined. In particular, we did not directly examine Piezo1-mediated Src spatial dynamics. Future studies addressing these mechanisms may provide additional insight into how mechanical cues are integrated into YAP-dependent signaling pathways.

From a translational standpoint, the association between Piezo1 activity, cytoskeletal organization, and nuclear signaling suggests that Piezo1 or its downstream effectors may represent potential targets for modulating mechanotransduction in cancer. However, further studies will be required to establish the functional consequences of these signaling changes in vivo.

This study has several limitations. First, the majority of mechanistic experiments were performed in MDA-MB-231 cells, and although comparative analyses with MCF-7 cells support our conclusions (Fig 2), validation in additional models will be necessary. Second, while Piezo1 activity was modulated genetically and pharmacologically, direct measurements of Piezo1-mediated calcium influx were not performed. Third, our conclusions are based on endpoint measurements of nuclear localization, and future studies using live-cell imaging will be required to resolve the dynamics of transcription factor trafficking. Finally, although our data support a cytoskeleton-associated mechanism, the precise molecular intermediates linking Piezo1 activity to cytoskeletal remodeling remain to be defined.

In conclusion, we identify Piezo1 as a mechanosensitive regulator that drives cytoskeletal remodeling and differentially regulates the nuclear localization of YAP and β-catenin. These findings provide a framework linking mechanical cues to cytoskeleton-dependent regulation of YAP and β-catenin localization (Fig 7).

## Acknowledgments

We thank J. Xu and B. Zhang at McMaster University for their valuable advice and support on this research.

## Data availability

All relevant data underlying the findings described in this manuscript are included in the article.

## References

1. Dupont S, Morsut L, Aragona M, Enzo E, Giulitti S, Cordenonsi M, et al. Role of YAP/TAZ in mechanotransduction. Nature. 2011;474: 179–184. doi:10.1038/nature10137

2. Harrison DL, Fang Y, Huang J. T-cell mechanobiology: Force sensation, potentiation, and translation. Frontiers in Physics. Frontiers Media S.A.; 2019. p. 45. doi:10.3389/fphy.2019.00045

3. Balkwill FR, Capasso M, Hagemann T. The tumor microenvironment at a glance. J Cell Sci. 2012;125: 5591–5596. doi:10.1242/JCS.116392

4. Acerbi I, Cassereau L, Dean I, Shi Q, Au A, Park C, et al. Human breast cancer invasion and aggression correlates with ECM stiffening and immune cell infiltration. Integrative Biology (United Kingdom). 2015;7: 1120–1134. doi:10.1039/c5ib00040h

5. Panciera T, Citron A, Di Biagio D, Battilana G, Gandin A, Giulitti S, et al. Reprogramming normal cells into tumour precursors requires ECM stiffness and oncogene-mediated changes of cell mechanical properties. Nat Mater. 2020;19: 797–806. doi:10.1038/s41563-020-0615-x

6. Venning FA, Wullkopf L, Erler JT. Targeting ECM disrupts cancer progression. Front Oncol. 2015;5: 165088. doi:10.3389/FONC.2015.00224/FULL

7. Piccolo S, Panciera T, Contessotto P, Cordenonsi M. YAP/TAZ as master regulators in cancer – modulation, function and therapeutic approaches. Nat Cancer. 2022;4: 9. doi:10.1038/S43018-022-00473-Z

8. Zanconato F, Cordenonsi M, Piccolo S. YAP/TAZ at the Roots of Cancer. Cancer Cell. Cell Press; 2016. pp. 783–803. doi:10.1016/j.ccell.2016.05.005

9. Wu Y, Su Z, Ge C, Barooj S, Hirota JA, Geng F. A stiffness-gated YAP-β-catenin axis orchestrates AXIN2 expression in metastatic breast cancer. iScience. 2026;29: 114405. doi:10.1016/j.isci.2025.114405

10. Dasgupta I, McCollum D. Control of cellular responses to mechanical cues through YAP/TAZ regulation. Journal of Biological Chemistry. 2019;294: 17693–17706. doi:10.1074/JBC.REV119.007963

11. Amit C, Padmanabhan P, Narayanan J. Deciphering the mechanoresponsive role of β-catenin in keratoconus epithelium. Scientific Reports. 2020;10: 21382-. doi:10.1038/s41598-020-77138-3

12. Wang X, Su L, Ou Q. Yes-associated protein promotes tumour development in luminal epithelial derived breast cancer. Eur J Cancer. 2012;48: 1227–1234. doi:10.1016/J.EJCA.2011.10.001

13. Li N, Lu N, Xie C. The Hippo and Wnt signalling pathways: crosstalk during neoplastic progression in gastrointestinal tissue. FEBS Journal. Blackwell Publishing Ltd; 2019. pp. 3745–3756. doi:10.1111/febs.15017

14. Reddy P, Deguchi M, Cheng Y, Hsueh AJW. Actin Cytoskeleton Regulates Hippo Signaling. PLoS One. 2013;8: e73763. doi:10.1371/JOURNAL.PONE.0073763

15. Akisaka T, Yoshida H, Inoue S, Shimizu K. Organization of Cytoskeletal F-Actin, G-Actin, and Gelsolin in the Adhesion Structures in Cultured Osteoclast. Journal of Bone and Mineral Research. 2001;16: 1248–1255. doi:10.1359/JBMR.2001.16.7.1248

16. Moleirinho S, Hoxha S, Mandati V, Curtale G, Troutman S, Ehmer U, et al. Regulation of localization and function of the transcriptional co-activator YAP by angiomotin. Elife. 2017;6: e23966. doi:10.7554/ELIFE.23966.001

17. Vanni G, Citron A, Suli A, Contessotto P, Caire R, Gandin A, et al. Microtubule architecture connects AMOT stability to YAP/TAZ mechanotransduction and Hippo signalling. Nature Cell Biology 2025;27: 1725–1738. doi:10.1038/s41556-025-01773-z

18. Cox CD, Bavi N, Martinac B. Biophysical Principles of Ion-Channel-Mediated Mechanosensory Transduction. Cell Rep. 2019;29: 1–12. doi:10.1016/j.celrep.2019.08.075

19. Raha A, Wu Y, Zhong L, Raveenthiran J, Hong M, Taiyab A, et al. Exploring Piezo1, Piezo2, and TMEM150C in human brain tissues and their correlation with brain biomechanical characteristics. Mol Brain. 2023;16: 1–11. doi:10.1186/S13041-023-01071-5

20. Pathak MM, Nourse JL, Tran T, Hwe J, Arulmoli J, Le DTT, et al. Stretch-activated ion channel Piezo1 directs lineage choice in human neural stem cells. Proc Natl Acad Sci U S A. 2014;111: 16148–16153. doi:10.1073/PNAS.1409802111

21. Deng J, Li Y, Zhang L, Liao W, Liu T, Shen F. Piezo1 regulates actin cytoskeleton remodeling to drive EMT in cervical cancer through the RhoA/ROCK1/PIP2 signaling pathway. Discover Oncology. 2025;16: 787. doi:10.1007/S12672-025-02474-7

22. Mousawi F, Peng H, Li J, Ponnambalam S, Roger S, Zhao H, et al. Chemical activation of the Piezo1 channel drives mesenchymal stem cell migration via inducing ATP release and activation of P2 receptor purinergic signaling. Stem Cells. 2020;38: 410–421. doi:10.1002/STEM.3114

23. Demagny J, Poirault-Chassac S, Ilsaint DN, Marchelli A, Gomila C, Ouled-Haddou H, et al. Role of the mechanotransductor PIEZO1 in megakaryocyte differentiation. J Cell Mol Med. 2024;28: e70055. doi:10.1111/JCMM.70055

24. Chen W, Bai Y, Patel C, Geng F. Autophagy promotes triple negative breast cancer metastasis via YAP nuclear localization. Biochem Biophys Res Commun. 2019;520: 263–268.

25. Chen W, Park S, Patel C, Bai Y, Henary K, Raha A, et al. The migration of metastatic breast cancer cells is regulated by matrix stiffness via YAP signalling. Heliyon. 2021;7: e06252. doi:10.1016/j.heliyon.2021.e06252

26. Konsavage WM, Yochum GS. Intersection of Hippo/YAP and Wnt/β-catenin signaling pathways. Acta Biochim Biophys Sin (Shanghai). 2013;45: 71–79. doi:10.1093/ABBS/GMS084

27. Gunst SJ, Zhang W. Actin cytoskeletal dynamics in smooth muscle: a new paradigm for the regulation of smooth muscle contraction. Am J Physiol Cell Physiol. 2008;295: C576. doi:10.1152/AJPCELL.00253.2008

28. Sugihara T, Werneburg NW, Hernandez MC, Yang L, Kabashima A, Hirsova P, et al. YAP Tyrosine Phosphorylation and Nuclear Localization in Cholangiocarcinoma Cells is Regulated by LCK and Independent of LATS Activity. Mol Cancer Res. 2018;16: 1556. doi:10.1158/1541-7786.MCR-18-0158

29. Hsu PC, Yang CT, Jablons DM, You L. The Crosstalk between Src and Hippo/YAP Signaling Pathways in Non-Small Cell Lung Cancer (NSCLC). Cancers (Basel). 2020;12: 1361. doi:10.3390/CANCERS12061361

30. Lamar JM, Xiao Y, Norton E, Jiang ZG, Gerhard GM, Kooner S, et al. SRC tyrosine kinase activates the YAP/TAZ axis and thereby drives tumor growth and metastasis. Journal of Biological Chemistry. 2019;294: 2302–2317. doi:10.1074/jbc.RA118.004364

31. Zhao B, Li L, Tumaneng K, Wang CY, Guan KL. A coordinated phosphorylation by Lats and CK1 regulates YAP stability through SCFβ-TRCP. Genes Dev. 2010;24: 72. doi:10.1101/GAD.1843810

32. Irtegun S, Wood RJ, Ormsby AR, Mulhern TD, Hatters DM. Tyrosine 416 Is Phosphorylated in the Closed, Repressed Conformation of c-Src. PLoS One. 2013;8: e71035. doi:10.1371/JOURNAL.PONE.0071035

33. Hu YJ, Wu X, Wang F, Jin Y, Jin Y, Liu Y, et al. Piezo1-mediated mechanotransduction controls osteocyte maturation and dendrite development via a YAP-CCN-Src signaling axis. Nature Communications. 2025;16: 10859-. doi:10.1038/s41467-025-65636-9

34. Hoa L, Kulaberoglu Y, Gundogdu R, Cook D, Mavis M, Gomez M, et al. The characterisation of LATS2 kinase regulation in Hippo-YAP signalling. Cell Signal. 2016;28: 488–497. doi:10.1016/J.CELLSIG.2016.02.012

35. Lei M, Wang W, Zhang H, Gong J, Wang Z, Cai H, et al. Cell-cell and cell-matrix adhesion regulated by Piezo1 is critical for stiffness-dependent DRG neuron aggregation. Cell Rep. 2023;42: 113522. doi:10.1016/J.CELREP.2023.113522

36. Zhou T, Gao B, Fan Y, Liu Y, Feng S, Cong Q, et al. Piezo1/2 mediate mechanotransduction essential for bone formation through concerted activation of NFAT-YAP1-ß-catenin. Elife. 2020;9. doi:10.7554/ELIFE.52779

37. Képiró M, Várkuti BH, Bodor A, Hegyi G, Drahos L, Kovács M, et al. Azidoblebbistatin, a photoreactive myosin inhibitor. Proc Natl Acad Sci U S A. 2012;109: 9402–9407. doi:10.1073/PNAS.1202786109

38. Zhao B, Li L, Lu Q, Wang LH, Liu CY, Lei Q, et al. Angiomotin is a novel Hippo pathway component that inhibits YAP oncoprotein. Genes Dev. 2011;25: 51–63. doi:10.1101/GAD.2000111

39. Hong W. Angiomotin’g YAP into the nucleus for cell proliferation and cancer development. Sci Signal. 2013;6. doi:10.1126/SCISIGNAL.2004573

40. Mana-Capelli S, Paramasivam M, Dutta S, McCollum D. Angiomotins link F-actin architecture to Hippo pathway signaling. Mol Biol Cell. 2014;25: 1676. doi:10.1091/MBC.E13-11-0701

41. Di X, Gao X, Peng L, Ai J, Jin X, Qi S, et al. Cellular mechanotransduction in health and diseases: from molecular mechanism to therapeutic targets. Signal Transduction and Targeted Therapy 2023;8: 1–32. doi:10.1038/s41392-023-01501-9

42. Orr AW, Helmke BP, Blackman BR, Schwartz MA. Mechanisms of mechanotransduction. Dev Cell. 2006;10: 11–20. doi:10.1016/j.devcel.2005.12.006

43. Eble JA, Niland S. The extracellular matrix in tumor progression and metastasis. Clinical & Experimental Metastasis 2019;36: 171–198. doi:10.1007/S10585-019-09966-1

44. Totaro A, Panciera T, Piccolo S. YAP/TAZ upstream signals and downstream responses. Nat Cell Biol. 2018;20: 888. doi:10.1038/S41556-018-0142-Z

45. Wang J, Jiang J, Yang X, Zhou G, Wang L, Xiao B. Tethering Piezo channels to the actin cytoskeleton for mechanogating via the cadherin-β-catenin mechanotransduction complex. Cell Rep. 2022;38: 110342. doi:10.1016/j.celrep.2022.110342

46. Cha B, Geng X, Mahamud MR, Fu J, Mukherjee A, Kim Y, et al. Mechanotransduction activates canonical Wnt/β-catenin signaling to promote lymphatic vascular patterning and the development of lymphatic and lymphovenous valves. Genes Dev. 2016;30: 1454–1469. doi:10.1101/GAD.282400.116

